# Cover crop residue diversity enhances microbial activity and biomass with additive effects on microbial structure

**DOI:** 10.1101/2021.02.08.430214

**Authors:** Xin Shu, Yiran Zou, Liz J. Shaw, Lindsay Todman, Mark Tibbett, Tom Sizmur

## Abstract

Cover crops have been widely used in agroecosystems to improve soil fertility and environmental sustainability. The decomposition of cover crop residues can have further effects on belowground communities and their activity, which is important for a series of soil functions (e.g., nutrient cycling and organic matter decomposition). We tested the effect of plant residues from a range of cover crop species on soil microbial activity and community assemblage. We predicted that cover crop residues would alter the soil microbial community and that a greater diversity of residues would enhance microbial decomposition. In an incubation study, we assessed the effect of crop residue diversity on microbial activity (soil respiration) and its consequent effects on microbial community composition (PLFA). We used either a biodiverse mixture of four cover crop residues (buckwheat, clover, sunflower and radish) or an equal mass of the residues of each of the individual species. The diverse mixture of cover crop residues had a significantly (*P* < 0.05) greater soil respiration rate, by 57.61 µg C g^−1^ h^−1^, than the average of the four individual residues, but did not have a significantly different soil microbial biomass or microbial community structure. This finding could be attributed to a greater diversity of organic resources increasing the number biochemical niches, and hence activating dormant microbial communities to increase microbial activity without affecting microbial biomass or community composition. Greater respiration from similar microbial biomasses suggests that microbial activity might be more efficient after a more diverse substrate input. This study confirms the positive impact of cover crop residues on soil microbial biomass and activity and highlights that mixtures of cover crop residues may deliver enhanced soil functions beyond the sum of individual cover crop residues.

## 1. Introduction

Integrating cover crops, non-cash crops grown primarily for the purpose of protecting or improving soil health between cash-crops in an arable crop rotation, is a popular tool to increase the sustainability of agricultural land management [1]. The use of cover crops has recorded benefits, such as increasing soil carbon storage, controlling soil erosion, enhancing agricultural productivity, reducing herbicide usage, managing diseases, increasing root biomass, and increasing the soil microbial biomass [1–4]. Since each individual species may excel at providing one or a few soil functions, a multi-species cover crop mixture is often used to maximise agroecological benefits [2]. For instance, mixtures of leguminous and non-leguminous species promote stable nitrogen (N) accumulation and mitigate N leaching, improving N capture, and thus providing a better green manure for the subsequent cash crop [1].

After the termination of cover crops, the decomposition of their residues can have further effects on belowground communities since the organic matter that is incorporated into soil is a significant driver of changes to the soil food web and microbial growth [5]. The biological degradation of plant residues which is one crucial part of soil functions, is mainly carried out by soil bacterial and fungal decomposers [6]. How quickly and efficiently microorganisms can degrade plant residues and assimilate the substrates into their biomass is closely linked to microbial stoichiometric requirements and the biochemical composition of plant residues [7]. In particular, the C/N ratio and the ratio of lignin to total N are the most important direct regulators of litter decomposition [8]. In N rich residues, decomposition can proceed without decomposers needing to scavenge exogenous N pools to meet stoichiometric requirements [6,9,10]. In contrast, N limitation could be a barrier for microorganisms to decompose residues with higher C/N ratio [6,9,10]. Different microbial groups may differ with respect to their preference in decomposing C substrates of distinct complexity and availability. For example, bacteria may be more effective competitors for the utilization of low molecular weight labile C substrates, while fungi may more easily mineralise larger and more complex biopolymers [11].

Due to differences in resource quality between plant species, the decomposition of residue mixtures can exhibit non-additive synergistic (mixture is decomposed faster than the predicted from single species), non-additive antagonistic (mixture is decomposed slower than the predicted) or additive effects (decomposition rate of mixture equals prediction) [12]. Gartner and Cardon (2004) reviewed 30 studies covering a wide range of ecosystems and found that about 67% of studies demonstrated a non-additive synergistic effect on plant residue decomposition of a mixture of plants, compared with what would have been predicted from single species. A possible explanation for the synergistic effect is nutrient transfer between plant residues, since nutrients released from rapidly decaying and higher quality plant residues (low C/N) can stimulate decay in lower quality plant residues (high C/N) [13]. Moreover, microbial N translocation between plant residues [14] and redistribution of N within fungal hyphal network [15] could mediate the decomposition of plant residue mixtures restricted by N-limitation. Therefore, mixing plant residues with a range of N concentrations could facilitate faster plant decomposition than the sum of the parts, producing a non-additive synergistic effect [16]. Mixing chemically contrasting plants provides a greater number of niches for microorganisms to exploit which allows functionally dissimilar microbial communities to coexist, and thus result in a greater microbial diversity and more efficient nutrient cycling [17,18]. Overall additive or antagonistic effects of plant residue mixtures on decomposition have also been reported. Decomposition rate of one plant residue could be suppressed by inhibitory compounds released by another plant residue. For example, inhibitory secondary compounds such as phenolics released from one plant residue can decrease decay rates of adjacent plant residues, and results in an overall reduced decomposition rate [13]. Competitive interactions between fungal communities could slow down the decomposition rate of litter mixtures, resulting in a non-additive antagonistic effect [19]. The factors responsible for whether a particular plant residue mixture induces an additive or non-additive (antagonistic or synergetic) effect are not all known, but could be related to, the chemical composition of the plant species selected, the mixing ratio, the number of species in the mix, and the soil function measured [12,20].

The benefits provided by a cover crop mixture over single species cover crop has been observed mostly in terms of its N retention in the soil and N supply to the following crop [1,21]. Many studies only focus on two species mixtures, particularly legume and non-legume [1]. However, more needs to know how cover crop mixture with diverse functional traits affects soil ecosystem services through underlying microbial communities [22]. We established a microcosm experiment to uncover the impacts of residues of cover crop mixtures comprising diverse functional traits on soil key function (respiration) and microbial community. We chose the residues of four individual cover crops, i.e., buckwheat, clover, radish sunflower, and a quaternary mixture which contains 25% residues from each individual by mass. These four plants are widely used as cover crops in agroecosystem because of their functional traits, i.e. buckwheat improves phosphorus efficiency, clover supplies N to subsequent cash crop, radish alleviates soil compaction, and sunflower provides considerable aboveground biomass to feed soil microorganisms after incorporation [22]. Therefore, the mixture of these four species may take maximum individual advantage and deliver better benefits. Moreover, these four plants are from different plant families (Polygonaceae, Fabaceae, Brassicaceae, and Asteraceae) and thus will synthesize family-specific primary and secondary metabolites which may have distinct impact on soil microbial communities and activities [23]. We hypothesized that 1) applying cover crop residues would increase soil respiration and microbial biomass due to the increased nutrient inputs; 2) soil microbial community structure would be shifted by cover crop residues in a species-specific way, and 3) the quaternary mixture which supplies a diverse array of resources may satisfy disparate microbial communities, facilitate microbial respiration, and thus exhibit a synergetic interaction, compared to the average of the four individual residue treatments.

## 2. Materials and methods

### 2.1 Soil samples and plant residues

Soil was sampled from an arable field on the University of Reading’s research farm at Sonning, Reading, UK (51.481152, −0.902188) in November 2018 after harvesting Spring barley (*Hordeum vulgare*). The soil is classified as silty loam Luvisol (World Reference Base classification) with pH (H_2_O) 6.3, 22.32 g C kg^−1^, 2.24 g N kg^−1^, 0.90 mg NH_4_^+^ −N kg^−1^, 2.75 mg NO_3_^−^ −N kg^−1^. Five surface soil samples (0-20 cm depth) were randomly sampled and mixed thoroughly to create one homogenous sample approximately 20 kg in weight.

Four cover crops, buckwheat (*Fagopyrum esculentum*), berseem clover (*Trifolium alexandrinum*), sunflower (*Helianthus annuus*), and oil radish (*Raphanus raphanistrum*), were grown between August and October 2018 in an area adjacent to where the soil was collected. Aboveground residues were harvested during the vegetative growth stage, dried at 70 °C and milled to pass through 0.05 mm mesh. The elemental composition of each cover crop residue, and the quaternary mixture (which consisted of 25% by mass of each of the four cover crops), is provided in Table 1 and S1.

**Table 1.** The nutrient contents of the cover crop residues used in experiments and the rate of C and N added to soils in the five treatments receiving cover crop residues.

| Element | Buckwheat | Clover | Radish | Sunflower | Mixture |
| --- | --- | --- | --- | --- | --- |
| Ca (mg kg<sup>-1</sup>) | 29600 | 21100 | 33400 | 35300 | 30200 |
| K (mg kg<sup>-1</sup>) | 21600 | 19800 | 35300 | 30700 | 26700 |
| Mg (mg kg<sup>-1</sup>) | 3830 | 2230 | 2080 | 7010 | 3720 |
| P (mg kg<sup>-1</sup>) | 4610 | 3860 | 5200 | 5310 | 4920 |
| S (mg kg<sup>-1</sup>) | 2420 | 2680 | 3790 | 4810 | 3780 |
| N (%) | 1.98 | 4.05 | 3.38 | 4.72 | 3.53 |
| C (%) | 38.8 | 42.1 | 38.9 | 39.4 | 39.8 |
| C/N ratio | 19.6 | 10.4 | 11.5 | 8.3 | 12.5 |
| Added C (mg C g<sup>-1</sup> soil) | 9.46 | 10.27 | 9.50 | 9.62 | 9.71 |
| Added N (mg N g<sup>-1</sup> soil) | 0.24 | 0.48 | 0.40 | 0.56 | 0.42 |

### 2.2 Soil incubation and measurement of respiration

Soil was sieved to pass 4 mm mesh and then pre-incubated for 7 days at 26 °C with a soil water content of 60% of the water holding capacity (0.22 g water g^−1^ soil). The treatments consisted of individual residue of buckwheat, clover, radish and sunflower, a quaternary mixture, and a control without any amendment with cover crop residues. For each treatment receiving plant residues, four replicate units were established by mixing 250 g of fresh soil (equivalent to 204.92 g dry soil) thoroughly with 5 g dry plant residues stored in a plastic bag. The amount of added C and N for each treatment was described in Table 1.

For each treatment and replicate, one aliquot of 200 g mixture of soil and plant was taken from the plastic bag, transferred, and loosely packed to a bulk density of 1 g cm^−3^ in a pot (200 ml). The pots were stored in a gas-tight plastic container (940 ml) with a headspace gas sampling port covered with a parafilm when not in sampling, and incubated at 26 °C. Soil respiration was measured as CO_2_ flux following [24] at 0, 1, 2, 3, 6, 7, 8, 9, 10, 13, 14, 30, 35, 42, 49, 63, 70, 77, 84 days after the addition of cover crop residues. Briefly, a parafilm was replaced with a Suba Seal® septa and 16 ml of headspace gas was collected by syringe after closure of the chamber for one hour, and stored in a pre-evacuated Labco exetainers® vials (12 ml). Gas from jars without soils was collected at the beginning of incubation to calibrate the background atmospheric CO_2_ concentration. Gas samples were analysed by gas chromatography with a thermal conductivity detector (Agilent GC 6890, UK). The universal gas law was used to determine the amount of CO_2_ as described by [24].

For each treatment and replicate, another three 5 g aliquots of the soils mixed with plant residues were stored in 30 ml polypropylene tubes and incubated at the same temperature as the samples for gas analysis for the analysis of PLFA at 1, 35 and 84 days after the addition of cover crop residues. Water was added to soils for gas and PLFA analysis regularly to compensate water loss.

### 2.3 PLFA

Soils were frozen on the day of sampling and then freeze-dried for downstream analysis of PLFA. PLFA was extracted following the method described by Sizmur et al., (2011) [25]. Briefly, 2 g of freeze-dried soil was extracted with 7.8 ml of one-phase extractant containing chloroform: methanol: citrate buffer (1:2:0.8 v/v/v). The extracted phospholipids were methanolized as fatty-acid methyl esters and then analysed using gas chromatography (Agilent Technologies 6890N, UK) [26]. Peaks were identified using a bacterial fatty acid methyl esters (BAME) mix (Sigma Aldrich, UK) and quantified using a 37-component fatty acid methyl esters (FAME) mix (Sigma Aldrich, UK). The biomass of each group of microorganisms was determined using the combined mass of fatty acids to which the group is attributed in Table S2.

### 2.4 Data analysis

All the statistics were conducted in R (version 3.5.2) [27] except for analysis of similarity (ANOSIM). We fit two separate linear mixed-effects model (LMM) using REML (restricted maximum likelihood) under the package “nmle” [28]. One LMM identifies the fixed effect from different treatment and time, and random effect from sample replicates on soil respiration rate and PLFA biomass. Another LMM identifies the fixed effect from the cover crop quaternary mixture and the average of the four individuals, and random effect from sample replicates on respiration rate and PLFA biomass.

Non-metric multidimensional scaling (NMDS) on Bray-Curtis distance of microbial communities (Hellinger transformed PLFA data) was performed using the “vegan” package [29] to distinguish soil microbial community structure influenced by cover crop residues. Bray-Curtis distance similarity matrices were calculated and used for a two-way analysis of similarity (ANOSIM) using Primer-e Version 7 (New Zealand) to test the significance of treatment and time on the soil microbial community structure.

## 3. Results

### 3.1 Effects of applying cover crop residues on soil respiration and microbial biomass

The addition of cover crop residues significantly (*P* < 0.001) increased soil respiration rate over 84 days, compared to the unamended control (Fig 1 and Table S3). Overall, the addition of buckwheat, clover, sunflower, radish, and a mixture containing 25% (w/w) of all four residues, resulted in an increase in soil respiration rate by 337, 340, 372, 373, and 359 µg C g^−1^ h^−1^, compared to the unamended control soil respiration rate of 29 µg C g^−1^ h^−1^ (Table S3). The response of soil respiration to cover crop residue addition was rapid but was followed by a steep decline in respiration rate over 84 days (Fig 1a). The respiration rate in the soil which received sunflower residue was significantly (*P* < 0.05) higher than buckwheat and clover which had the lowest respiration rate (Table S4).

**Fig 1.**
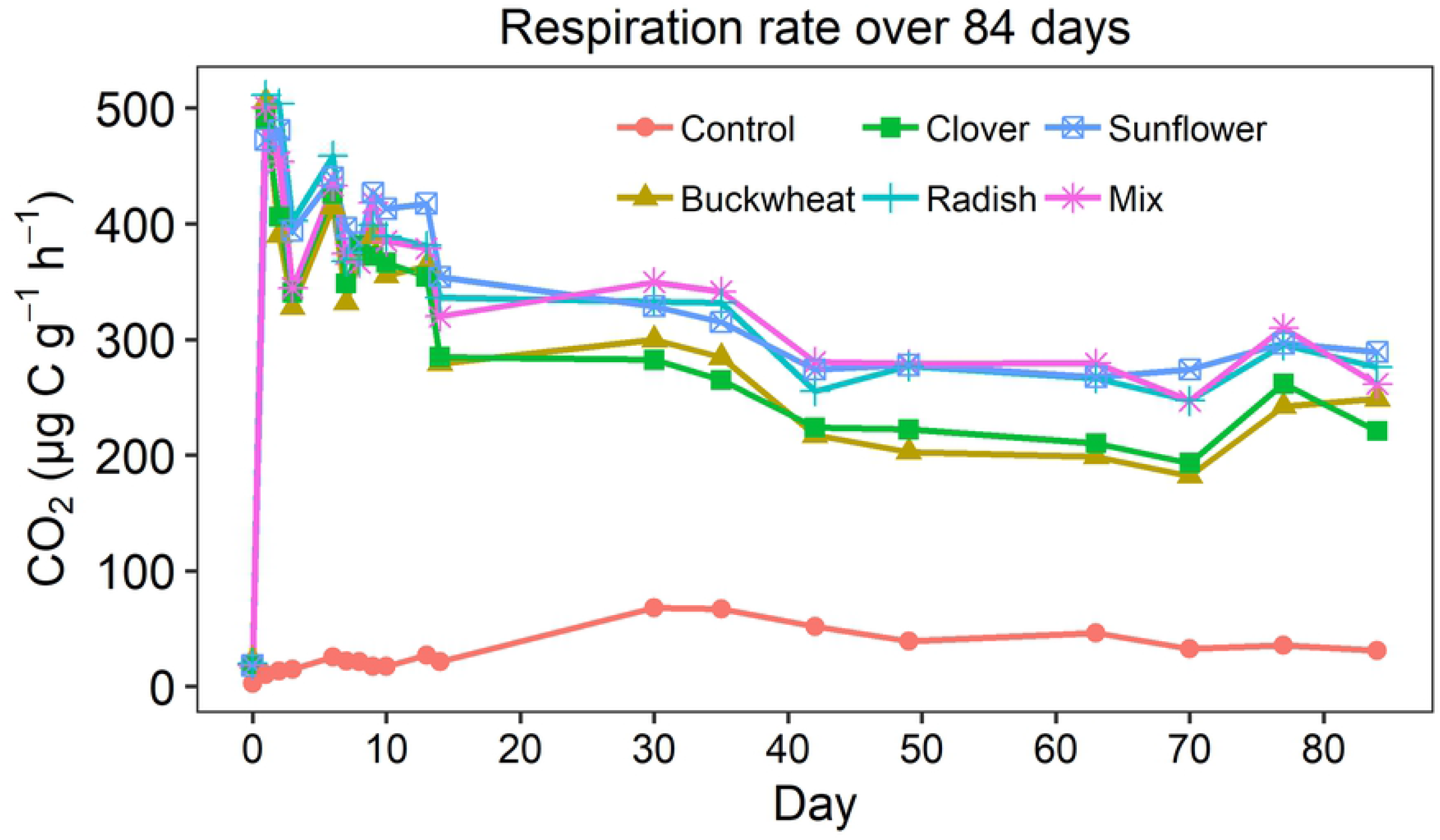
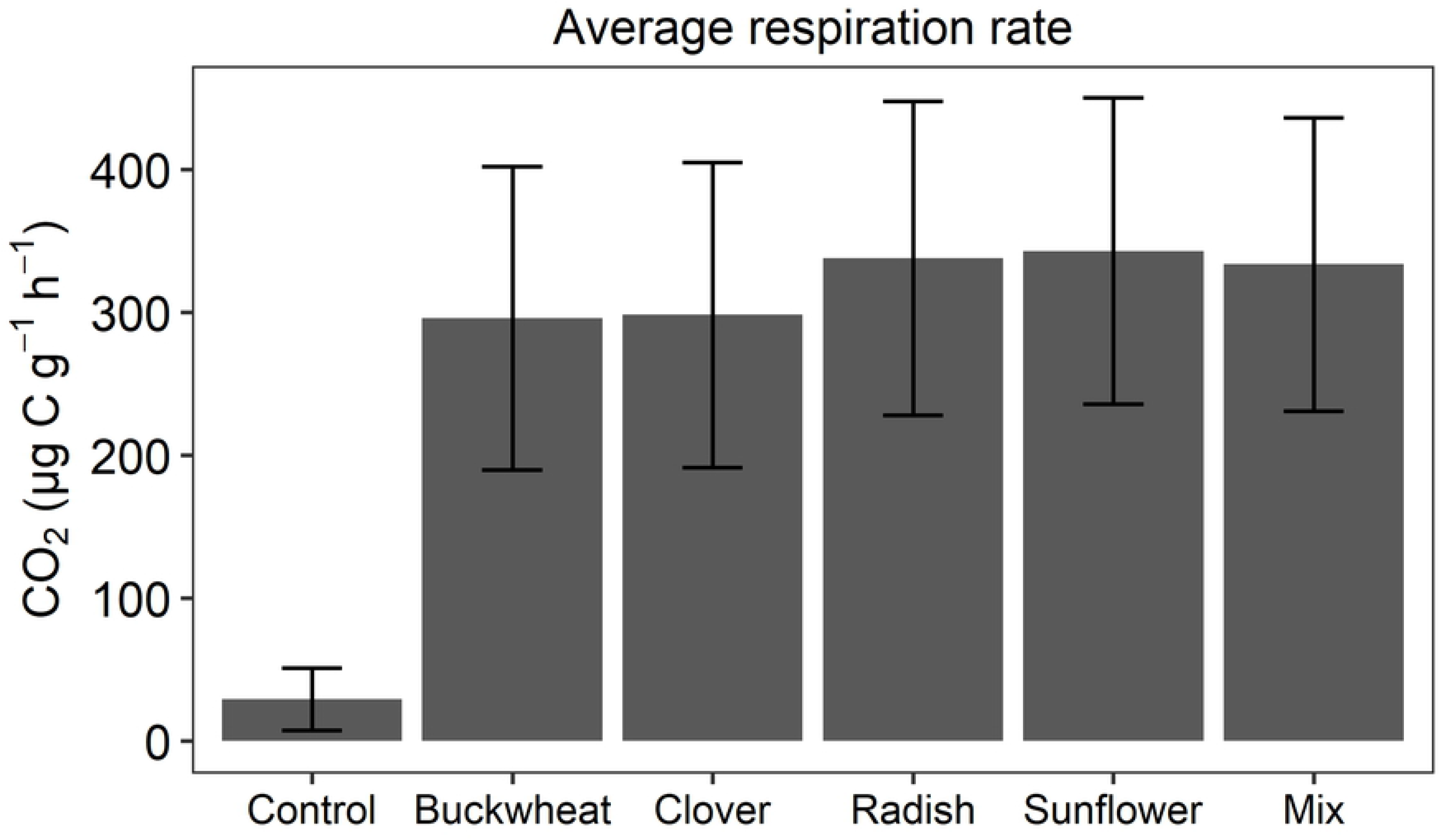
Soil respiration rate in different treatments over 84 days (a), and the average respiration rate (b). Mix is the quaternary mixture of buckwheat, clover, radish and sunflower containing 25% (w/w) of all four residues. Control is soil without the addition of plant residues.

Cover crop residues also significantly (*P* < 0.001) increased the growth of microorganisms, as revealed by a greater total PLFA biomass, compared to the unamended control treatment (Fig 2 and Table S3). Overall, the addition of cover crop residues resulted in an increase of total PLFA biomass by 46, 64, 53, 66, and 59 µg C g^−1^ soil for buckwheat, clover, sunflower, radish, and the quaternary mixture, compared to the unamended control of which total PLFA was 18 µg C g^−1^ (Table S3). There were no significant differences in total PLFA biomass between the treatments receiving different cover crop residues (Table S4).

**Fig 2.**
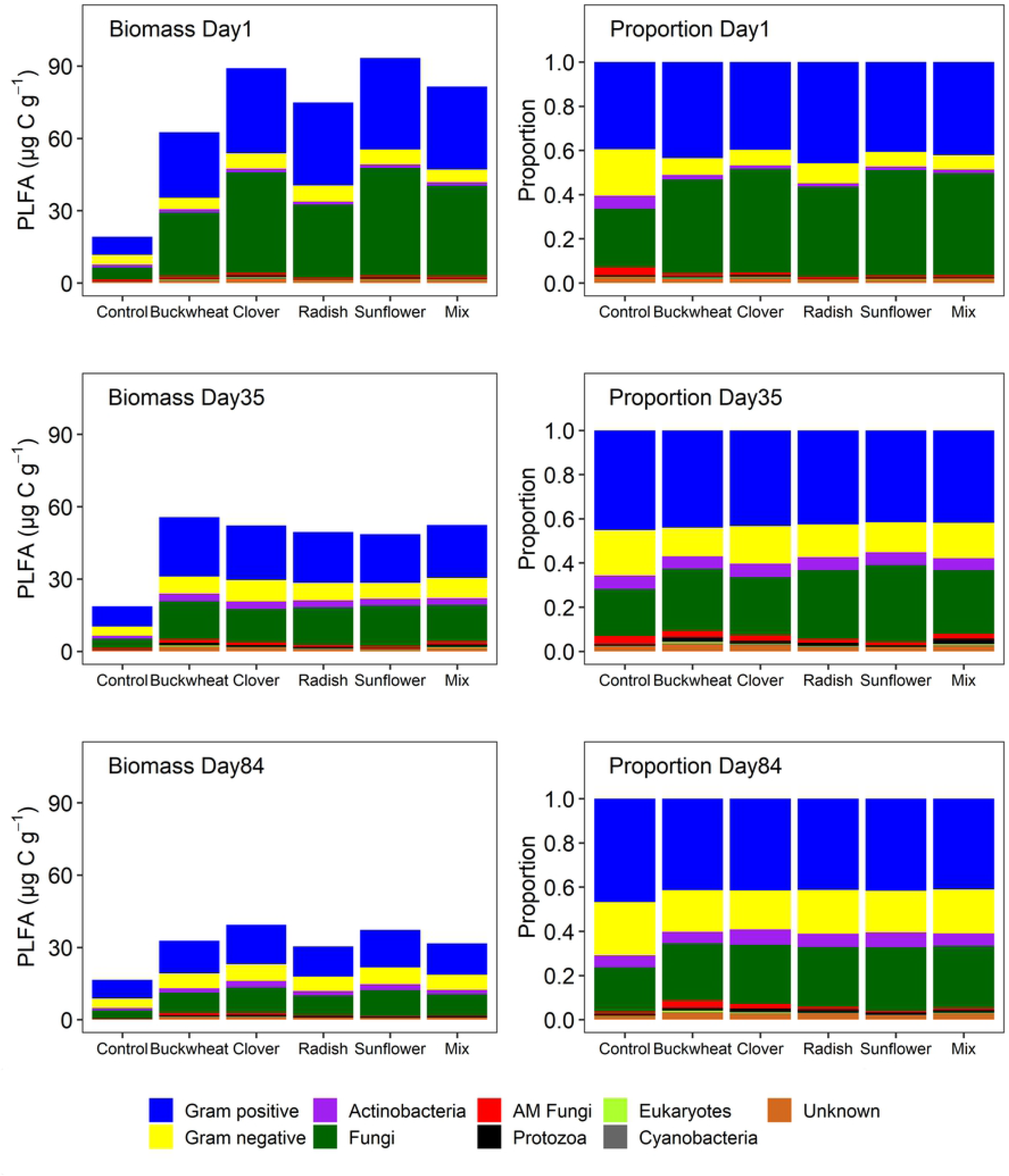
PLFA biomass (µg C g^−1^) and proportion in all treatments at day 1, 35 and 84. Mix is the quaternary mixture of buckwheat, clover, radish, and sunflower containing 25% (w/w) of all four residues. Control is soil without any addition. AM fungi is arbuscular mycorrhiza fungi. Fungi or protozoa are not included in eukaryotes, and *actinobacteria* are not included in the Gram-positive bacteria.

The biomass of fungi (*P* < 0.001), Gram-positive bacteria (*P* < 0.001), Gram-negative bacteria (*P* < 0.001), *cyanobacteria* (*P* < 0.05) and a group of fatty acids that cannot be attributed to an individual microbial group (*P* < 0.05) were significantly increased by the addition of cover crop residues (Table S3 and Fig 2). Arbuscular mycorrhizal fungi and *actinobacteria* onlyn occupy 0.5-3% and 1.5-6.9% of the total microbial biomass, respectively (Fig 2) and did not significantly respond to any cover crop residue addition (Table S3). The abundance of fatty acids attributed to protozoa (20:3 ω6c and 20:4 ω6c) and a general fatty acid biomarker for eukaryotes (18:3 ω6) were significantly (*P* < 0.05) increased by the addition of buckwheat, clover and the quaternary mixture of residues (Fig 2 and Table S3). Except for radish, the application of cover crop residues significantly (*P* < 0.05) increased the abundance of a general fatty acid biomarker for eukaryote.

The ratio of fungi to bacteria in the unamended soil is 0.35. On average, the ratio of fungi to bacteria was significantly (*P* < 0.001) increased by 0.35, 0.46, 0.54, 0.30 and 0.44 following the addition of buckwheat, clover, sunflower, radish, and the quaternary mixture of residues (Table S3 and Fig S1).

### 3.2 Changes of microbial community structure under different crop residues over time

Adding cover crop residues significantly (*P* < 0.001) altered the soil microbial community structure regardless of the species of cover crop (Fig 2 and 3 and Table 2). Compared to the unamended control, the soil microbial communities in the cover crop residue treatments had a significantly (*P* < 0.001) greater relative abundance of fungi, and a significantly (*P* < 0.001) lower relative abundance of Gram-negative bacteria, arbuscular mycorrhizal fungi and *actinobacteria* (Table S5). Incorporation of buckwheat, radish, and the quaternary mixture significantly (*P* < 0.05) increased the relative abundance of Gram-positive bacteria, whereas no significant effects were found due to the addition of sunflower or clover residues. Focussing just on the treatments receiving cover crop residues revealed that soil microbial community composition was significantly different (*P* < 0.001) for all pairwise comparisons (Table 2). The residues of buckwheat or sunflower led to the development of microbial communities with the greatest dissimilarity between each other than any other comparison in this experiment. A significant (*P* < 0.001) difference in microbial community composition was found between the three time points, with a greater difference between day 1 and day 35 or 84 than between day 35 and day 84 (Table 2).

**Fig 3.**
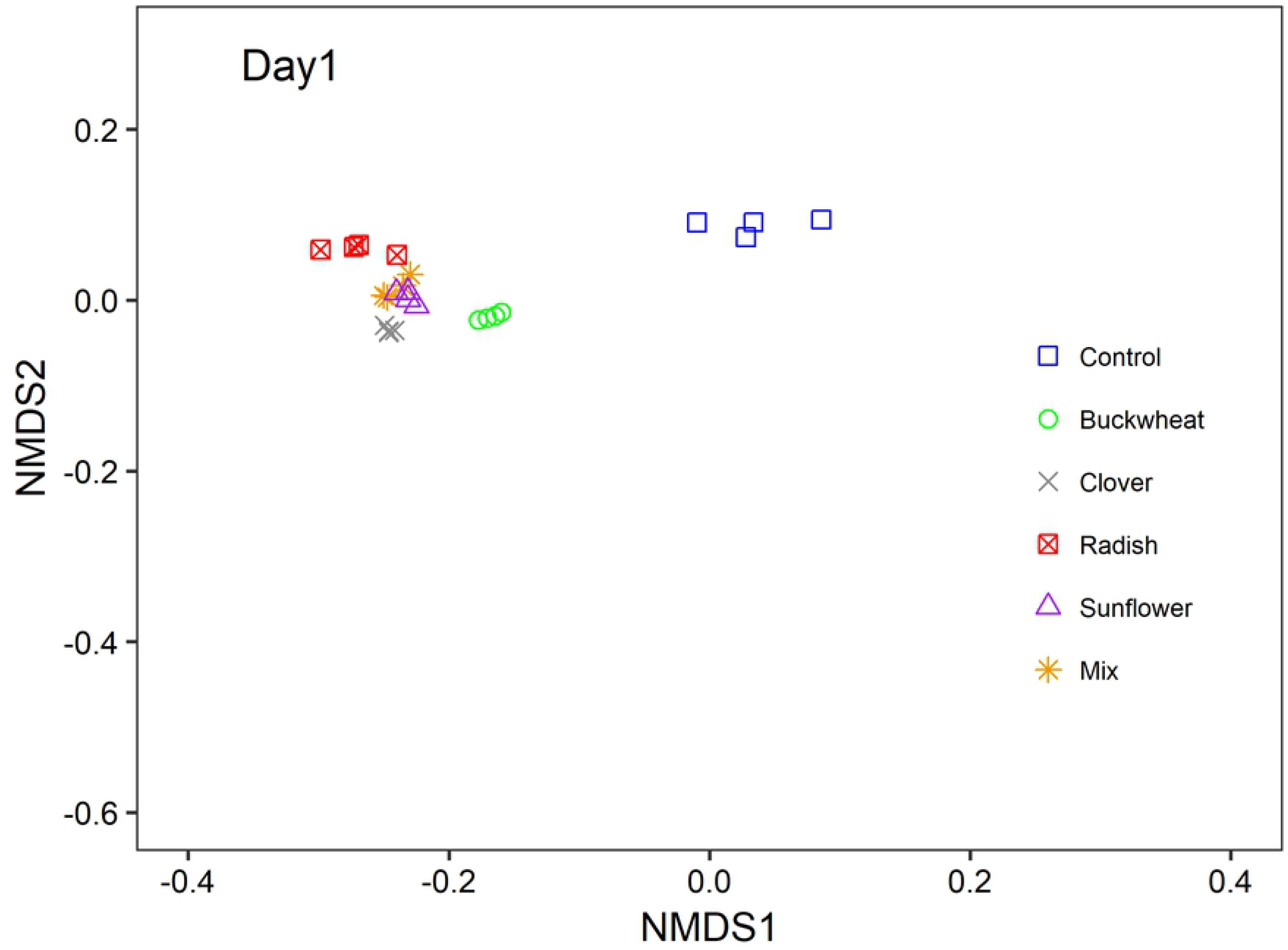
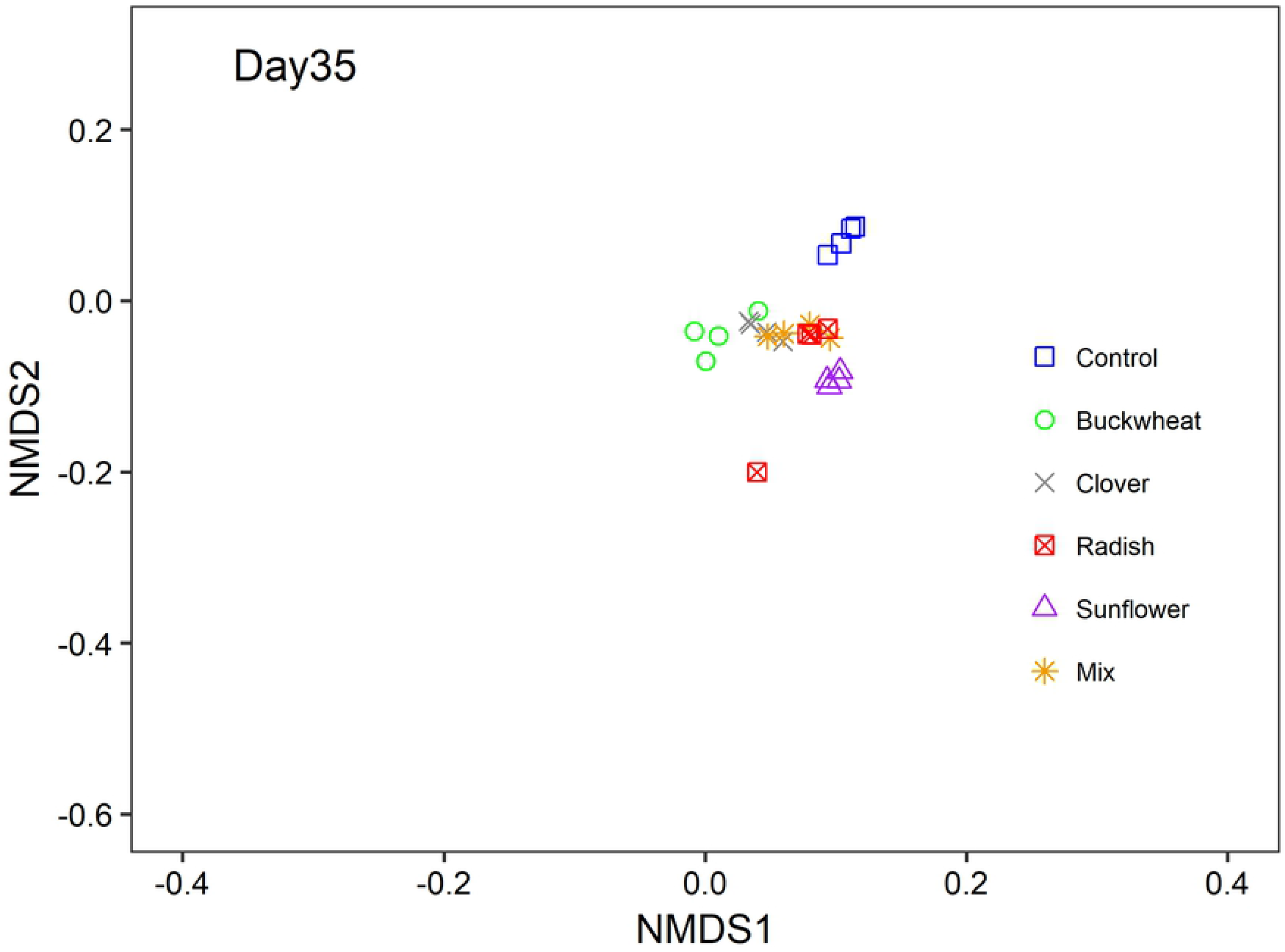
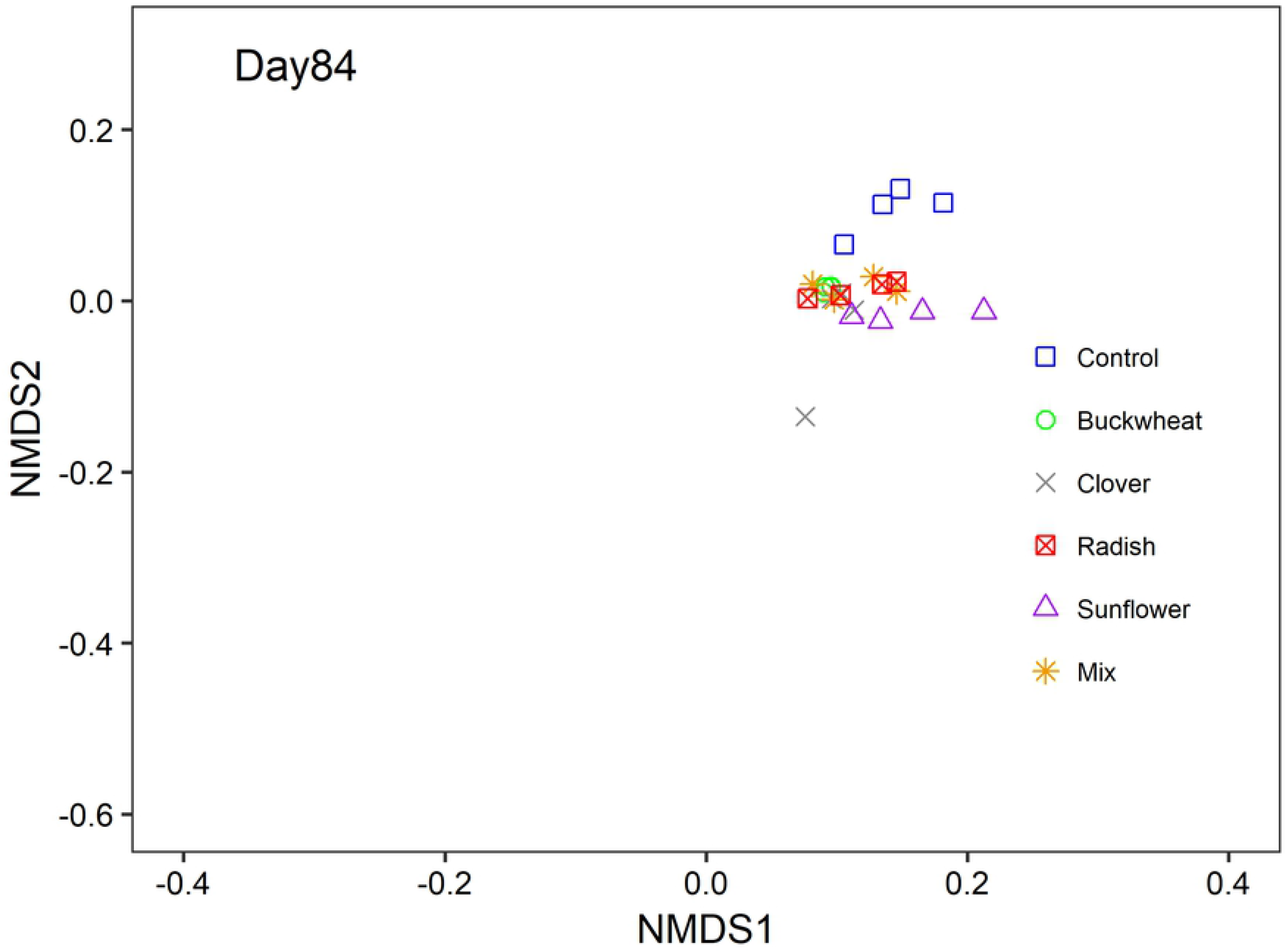
Microbial community structure in Day 1 (a), Day 35 (b), Day 84 (c). Non-metric multidimensional scaling (NMDS) based on Bray-Curtis distance analysis of Hellinger transformed abundance of phospholipid fatty acid (PLFA) biomarkers. Dots are samples. Stress = 0.0929, R^2^ = 0.991.

**Table 2.** ANOSIM results (R value and P value) of pairwise comparisons of the soil microbial community composition (PLFA profile) between different treatments or different time points after applying cover crops.

|  | R value | P value |
| --- | --- | --- |
| Control/ Mix | 0.979 | < 0.001 |
| Control/ Radish | 0.844 | < 0.001 |
| Control/ Sunflower | 0.990 | < 0.001 |
| Control/Buckwheat | 0.972 | < 0.001 |
| Control/Clover | 0.903 | < 0.001 |
| Mix/ Radish | 0.285 | < 0.001 |
| Mix/ Sunflower | 0.785 | < 0.001 |
| Mix/Buckwheat | 0.847 | < 0.001 |
| Mix/Clover | 0.688 | < 0.001 |
| Buckwheat/ Clover | 0.712 | < 0.001 |
| Buckwheat/ Radish | 0.743 | < 0.001 |
| Buckwheat/ Sunflower | 0.944 | < 0.001 |
| Clover/ Radish | 0.628 | < 0.001 |
| Clover/ Sunflower | 0.826 | < 0.001 |
| Radish/ Sunflower | 0.618 | < 0.001 |
| Day 1/Day 35 | 0.939 | < 0.001 |
| Day 1/ Day 84 | 0.977 | < 0.001 |
| Day 35/ Day 84 | 0.740 | < 0.001 |

### 3.3 Difference between mixture and the average of four individual cover crop residues

Compared to the average of the four individual cover crop residue treatments, the soil respiration rate was not significantly higher in the quaternary mixture treatment when all measurements are considered (Fig 4 and Table S6). However, during the period 30 to 84 days after adding cover crop residues, the soil respiration rate in the mixture treatment was significantly (*P* < 0.05) greater by 57.61 µg C g^−1^ h^−1^ than the average of four individual residues (Fig 4 and Table S6).

**Fig 4.**
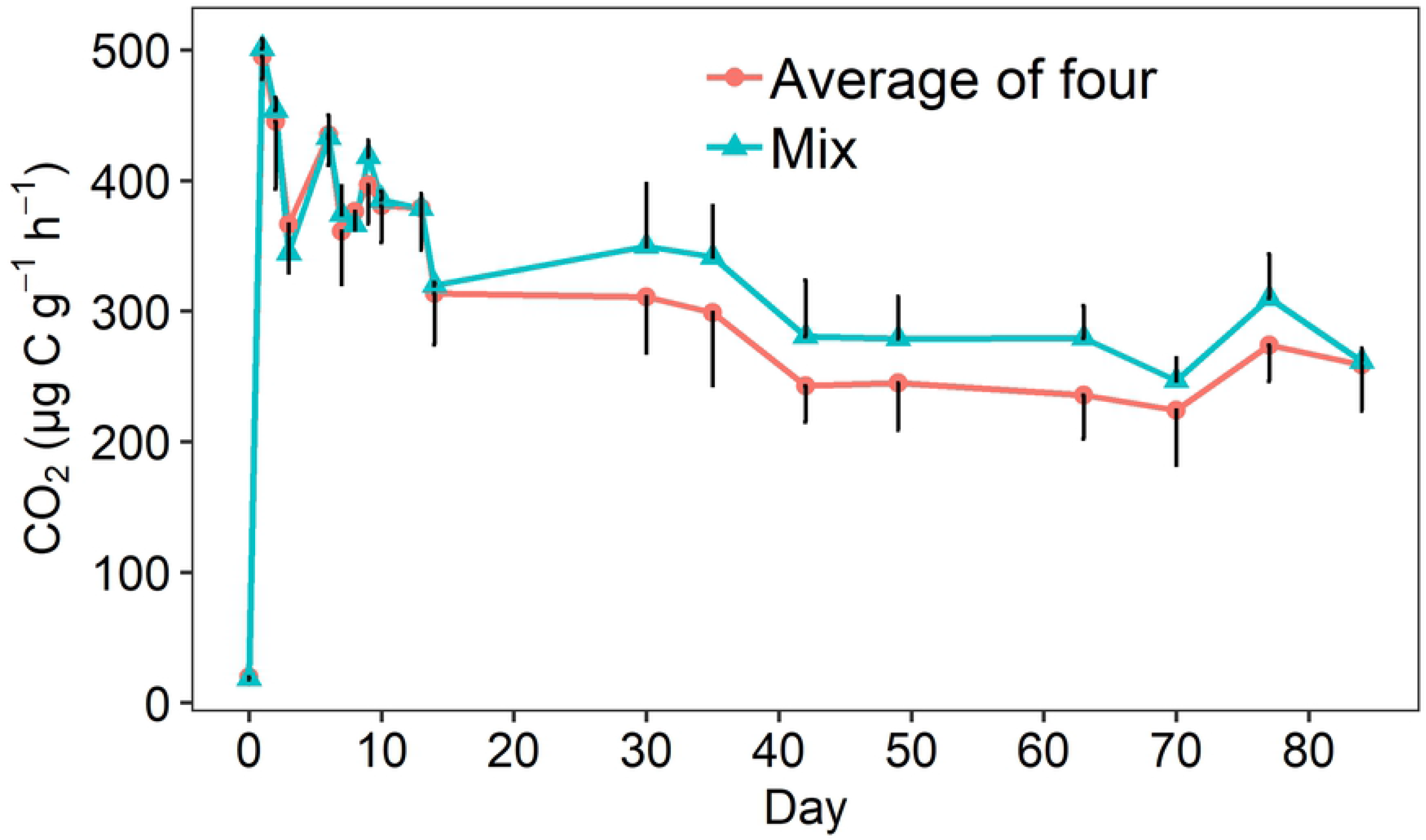
The comparison of soil respiration rate between soils amended with the cover crop residue mixture and the average of four treatments amended with the residues of individual cover crop species. Mean and error bar (standard deviation). To aid visualization, the error bar for mixture is positive whereas for the average of four is negative.

The mixture treatment had a slightly, but not statistically significant, greater total microbial biomass than the average of the four individuals at 1 and 35 days after cover crop residue application, but a lower soil microbial biomass after 84 days (Fig 5). The biomass of fungi and Gram-positive bacteria were slightly greater in the quaternary mixture treatment than the average of the four individual cover crop treatments one day after residue incorporation (Fig 5), but there was no overall significant difference between the mixture treatment and the average of four individual treatments across the three time points (Table S6). Although different microbial biomarkers responded to the mixture differently, none of these differences were significant (Fig S2).

**Fig 5.**
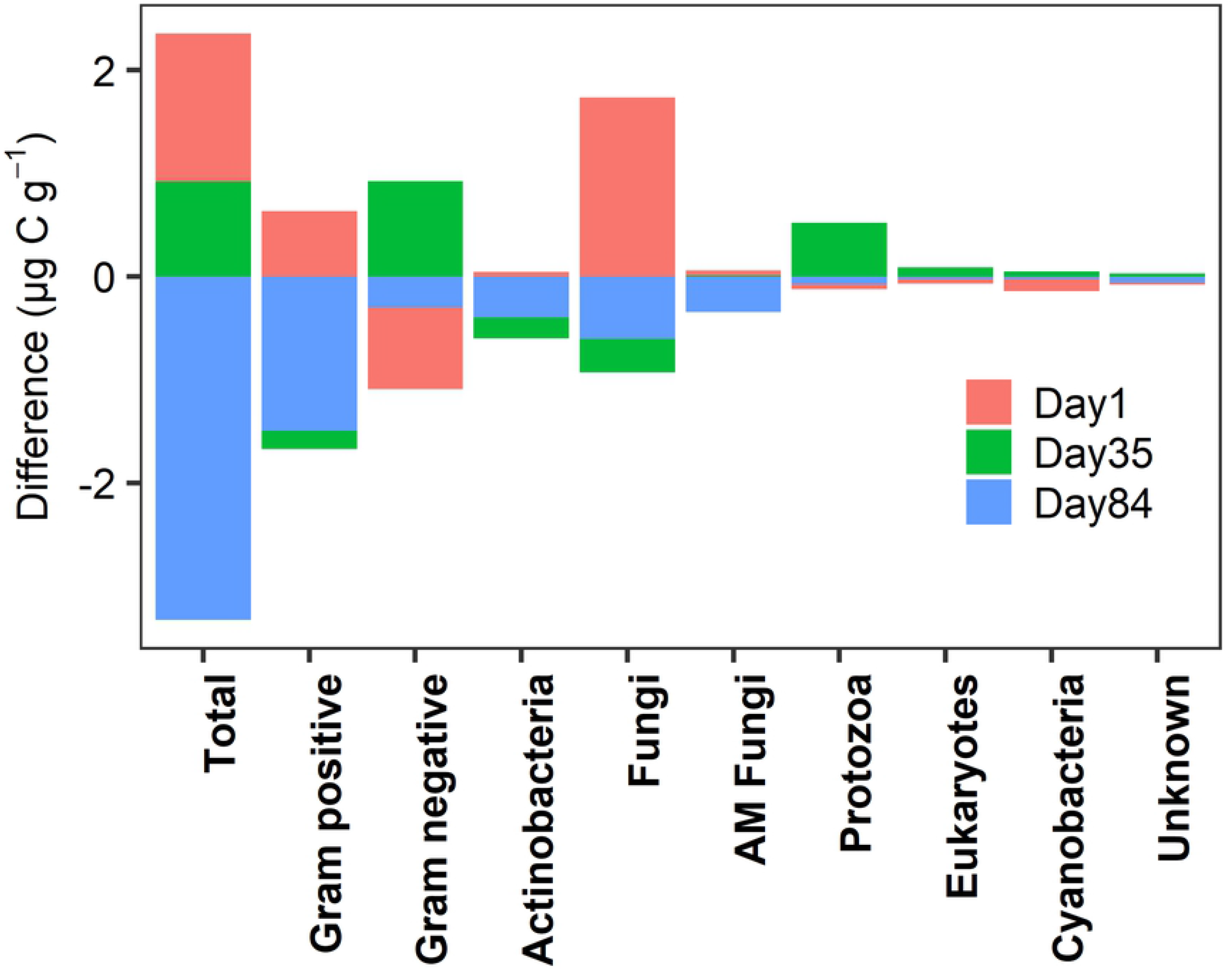
Mass difference of microbial groups between soils amended with the cover crop residue mixture and the average of four treatments amended with the residues of individual cover crop species over 84 days. Positive value means mixture has greater biomass than the average of the four individuals, and vice versa.

## 4. Discussion

### 4.1 Cover crop residues increased soil respiration and microbial biomass

The rapid elevation in respiration rate following a one-time addition of cover crop residues (Fig 1), compared to unamended soil, highlights an intensive decomposition period when the most labile components of plant residues were mineralised [30]. The highest respiration rate was found in the treatment with sunflower which also had the lowest C/N ratio, confirming low C/N ratio plant residues can be more easily mineralised by soil microorganisms [8]. When plant residues are decomposed, available nutrients and organic compounds can be quickly assimilated into microbial biomass [5]. This is why we observed a large increase of PLFA biomass in all the treatments receiving cover crop residues.

The increased fungi to bacteria ratio stimulated at the beginning of the degradation process implies fungi have greater C use efficiency and faster exploitation of fresh substrates than bacteria, particularly Gram-negative bacteria [31]. This is similar to previous work which found a relatively greater increase in fast-growing fungi that specialise in metabolising C-rich material than bacteria [32]. Fungi and bacteria apply different feeding strategies. Bacteria are more competitive in decomposing simple carbohydrates (e.g. glucose) than fungi [11], while the growth of fungi can be favoured by litters containing O-alkyl C groups in biopolymers such as polysaccharides and polypeptides [5]. To capture more essential resources and occupy a greater niche, fungi may apply antagonistic strategies by producing secondary metabolites (e.g. tolaasin) to inhibit the growth of bacteria [33]. With the help of hyphae, fungi may gain access to less accessible substrates, such as cellulose fibrils, which are unlikely to be decomposed by bacteria [33]. In fact, a fungi dominant community is more efficient at recycling energy from decomposing complex organic compounds than bacteria, and thus contributes to a soil ecosystem with greater microbial biomass [7].

### 4.2 Cover crop residues changed soil microbial biomass and community

Our study revealed a significant (*P* < 0.001) separation between the soil microbial community structure of treatments which received cover crop residues and the unamended control treatment (Table 2 and Fig 3). This observation is consistent with a previous study that demonstrated that the incorporation of plant residues provides a rich resource to boost the growth of microorganisms and alter microbial community structure [34]. In such a resource rich system, competitors which excel at maximising resource capture will reproduce faster and occupy more niches [35]. It is unsurprising that AM fungi biomass did not increase in our experiment, given the absence of living potential host plants. The relatively greater proportions of fungi and Gram-positive bacteria were mirrored by a decrease in the relative proportion of Gram-negative bacteria and *Actinobacteria* (in addition to AM fungi), suggesting saprotrophic fungi and Gram-positive bacteria are more efficient in assimilating available resources to prevail and survive under the same environment. Different cover crop residues induced different microbial community structures. The largest distinction was found between buckwheat and sunflower which is consistent with them having the biggest difference in C/N ratio, confirming that plant-derived substrate biochemical composition selects microbial community composition [34].

Soil microbial community composition is a result of complex interactions between plant residues, soil physicochemical properties, the surrounding environment, and time. By the end of the incubation, the difference between the soil microbial community in the control treatment and the treatments where cover crop residues were applied was smaller than the difference just one day after application. This return indicates a shift from a stochastic stage where residue inputs drive succession, to a deterministic phase where soil environmental factors play major roles in microbial succession [36]. Moreover, with the decomposition of plant residues, the life history strategy of the microbial communities may change from assimilating resources into biomass to the resource acquisition strategy in which more energy will be spent on producing extracellular enzymes to decompose complex resources rather than community succession or evolution [35].

### 4.3 The mixture delivered different effects on microbial respiration and community

Our results found no significant overall difference in CO_2_ flux between the quaternary mixture of cover crop residues and the average of the four individual residues over 84 days of incubation (Fig 4), indicating that a mixture of crop residues results in an overall additive effect on crop residue degradation over time. This is consistent with a previous study that examined the decomposition of litters which contrasted in their specific rate of decomposition but decomposed at a rate that was additive when in a mixture [37]. The C/N ratios of plant residues in our study are all less than 25, indicating that N should not limit microbial decomposition of plant residues [38]. Since the amount of added C is the same between the mixture and the average of the four individual residues (both are 9.71 mg C g^−1^ soil), it is not surprising that the decomposition rate in the mixture is similar to the average of four individual species.

Our study found a synergistic effect of the quaternary mixture of cover crop residues compared to the average of the four individual cover crop residues on soil respiration between 30 and 84 days after residue application. Our results are consistent with previous studies reporting that synergistic effects become stronger and more frequent as decomposition proceeds, which suggests that incubation time is an important factor to capture the additive or non-additive effect from the mixture [20,39]. Both antagonistic and synergistic effects could occur because the presence of certain compounds within one plant residue that promotes or inhibits the decomposition of other parts of a plant residue mixture [15]. For instance, the presence of phenolics and tannins could suppress microbial resource assimilation and slow down litter decomposition [13]. Consequently, the trade-off between this inhibition or promotion determines the overall rate of decomposition [20]. And the relative balance of antagonistic and synergistic effect in the mixture could result in an overall additive effect [20]. As resource and microbial community composition shifts as decomposition progresses, stoichiometric requirements for microbial catabolic and anabolic process can also shift, and potentially influence microbial C use efficiency with consequences for the amount of respired CO_2_ [7].

Although each plant residue induced a distinct microbial community structure, the incorporation of a mixture of cover crop residues did not generate major or statistically significant changes in the composition of the soil microbial community, compared to the average of the four communities in the soils which received individual cover crop residues (Fig 3). This suggests while plant residue identity is an important driver of soil microbial community, the diversity of plant residues per se might not be [40]. One reason could be limited communication between microbial “hotspots” associated with individual plant residues due to spatial isolation of these “hotspots”. Heterogenous pore space in the soil system limits the diffusion of nutrients and water between different “hotspots”, and thus weakens the coevolutionary relationship between microbial communities [41]. Additionally, the overarching role of soil environmental conditions might override the effects of plant residue diversity on microbial community [37].

We did not observe a synergistic effect of the cover crop residue mixture on the soil microbial community during the period 30 to 84 days when a synergistic effect on soil respiration was observed, which could be attributed to microbial C allocation strategy. Microorganisms allocate C to form extracellular enzymes, polysaccharides and other polymeric substances, and adjust osmolytes to combat environment changes and increase energy use efficiency, which may not necessarily change their community profile [42]. A recent study also found that a mixture of plant residues did not change microbial biomass but did increase microbial activity, compared to monocultures, indicating a strong phenotypic plasticity in the resident microbial communities [43]. A diverse array of substrates may enhance synergistic microbial interactions between the resident communities. For example, fungi may facilitate the penetration of bacteria into a plant residue-associated “hotspot” where both can degrade polymer and accelerate plant residue decomposition, which may not change microbial cell membrane composition, as measured by the PLFA method [15]. Increasing the diversity of substrates available to the soil microbial communities may activate the metabolism of dormant microbial communities [44]. These physiological changes in the soil microbial community, such as the transformation of dormant microorganisms to an active state and subsequent enhanced microbial connection/interaction, may not be expressed by differences in PLFA profiles. This study cannot rule out that the synergistic effect of the residue mixture on respiration may come as a result of an enhanced “priming effect” on the decomposition of native soil organic matter and necromass. It is reported that a sunflower-wheat mixture could induce a greater positive priming effect than the sunflower alone [45]. It is a possibility that interspecific interaction between extracellular enzymes produced by microbial communities to depolymerise different components of the mixture may also depolymerise native organic matter and thus enhance microbial metabolism and stimulate CO_2_ flux. Future research applying isotopic techniques to trace the fate of added C, investigating the metabolically active microbial communities and determining the interconnectivity of soil microbial networks, will help us to further understand reasons for the observed synergistic effects of cover crop residue mixtures on soil respiration. The effects of crop residue mixtures on the overall decomposition rate are consequences of interactions between different plant species, shifts in microbial life history strategy, and resource flow in the soil ecosystem.

## 5. Conclusion

Our results demonstrated cover crop residues significantly (*P* < 0.001) increased microbial biomass and soil respiration and altered the soil microbial community structure. Fungi and Gram-positive bacteria were the most responsive microbial groups to plant residue additions. The mixture of cover crop residues (comprising 25% by mass of each of the four individual residues) significantly (*P* < 0.05) increased soil respiration rate during the period 30 to 84 days after residue incorporation, compared to the average of the four individual residues, but there was no significant difference in soil microbial biomass or composition observed. Non-additive effects are strongly influenced by time, highlighting the importance of microbial life history strategies after exposure to crop residue mixtures. This study suggests that mixtures of cover crops could enhance microbial activity beyond that observed in soils receiving the residues of individual cover crop residues without altering soil microbial community composition.

## Acknowledgements

The authors wish to thank Kings Crops, a division of Frontier Agriculture Ltd, for supplying cover crop seed. We acknowledge the assistance of Anne Dudley, Karen Gutteridge, Fengjuan Xiao, Marta O’Brian, Sean Coole with laboratory analysis, and Dedy Antony, Alex Adetunji Adekanmbi and Chinonso Chukwuma Ogbuagu with soil sampling.

---

**Fig S1 The ratio of fungi to bacteria in all the treatments. Means and standard deviation (error bars)**.

**Fig S2 Mass difference of all measured biomarkers between the mixture and the average of the four individuals (buckwheat, clover, radish and sunflower). Positive value means biomarker’s biomass is greater in the mixture than the average of the four individuals, and vice versa**.

**Fig S3 The scores of non-metric multidimensional scaling (NMDS) based on Bray-Curtis distance analysis of Hellinger transformed abundance of phospholipid fatty acid (PLFA) biomarkers**.

**Table S1 The trace element contents of cover crop residues used in the experiment**

**Table S2 The attribution of PLFA biomarkers to microbial groups**

**Table S3 Coefficients of linear mixed effects model analysing the fixed effects from different treatments and day as well as their interaction on soil respiration rate (µg C g**^**-1**^ **h**^**-1**^**), fungi to bacteria ratio, and the biomass (µg C g**^**-1**^**) of total PLFA, fungi, Gram-positive bacteria, Gram-negative bacteria, eukaryote, protozoa, AM fungi, actinobacteria, cyanobacteria, and unknown microbial group. DF means degree of freedom. For all microbial group, the DF are the same as total PLFA. The effects are the effects of different treatments with a comparison to the control which received no cover crop residues. *, **, and *** means significant difference at *P* < 0.05, < 0.01, and <0.001. Abbreviations are F/B ratio (fungi to bacteria ratio), G+ bacteria (Gram-positive bacteria), G-bacteria (Gram-negative bacteria), AMF (arbuscular mycorrhiza fungi)**.

**Table S4 Multiple pairwise comparison of the effect between different plant residues on respiration rate (µg C g**^**-1**^ **h**^**-1**^**), fungi to bacteria ratio, and the biomass (µg C g**^**-1**^**) of total PLFA, fungi, Gram-positive bacteria, Gram-negative bacteria, eukaryote, protozoa, AM fungi, actinobacteria, cyanobacteria, and unknown microbial group. *, **, and *** means significant difference at *P* < 0.05, < 0.01, and <0.001. Abbreviations are F/B ratio (fungi to bacteria ratio), G+ bacteria (Gram-positive bacteria), G-bacteria (Gram-negative bacteria), AMF (arbuscular mycorrhiza fungi)**.

**Table S5 Coefficients of linear mixed effects model analysing the fixed effects from different treatments and day as well as their interaction on relative abundance (proportion) of fungi, Gram-positive bacteria, Gram-negative bacteria, eukaryote, protozoa, AM fungi, actinobacteria, cyanobacteria, and unknown microbial group. For all microbial group, the DF (degree of freedom) are the same. The effects are the effects of different treatments with a comparison to the control which received no cover crop residues. *, **, and *** means significant difference at *P* < 0.05, < 0.01, and <0.001**.

**Table S6 Coefficients of linear mixed effects model analysing the fixed effects from mixture plant residues and day as well as their interaction on soil respiration rate (µg C g**^**-1**^ **h**^**-1**^**), fungi to bacteria ratio, and the biomass (µg C g**^**-1**^**)of total PLFA, fungi, Gram-positive bacteria, Gram-negative bacteria, eukaryote, protozoa, AM fungi, actinobacteria, cyanobacteria, and unknown microbial group. DF means degree of freedom. The effects are the effects of the quaternary mixture of cover crop residues with a comparison to the average of the four individuals. The respiration was compared at two time scales: over 84 days or from day 30 to 84 days. *, **, *** indicates significant at *P* < 0.05, 0.01, and 0.001**.

